# Promiscuous cytochrome P450s confer metabolic resistance to synthetic auxin herbicides in the weedy grass *Echinochloa phyllopogon*

**DOI:** 10.1101/2025.07.29.667533

**Authors:** Pattarasuda Chayapakdee, Kowit Hengphasatporn, Takuya Yamaguchi, Niña Gracel Dimaano, Yasuteru Shigeta, Yukari Sunohara, Hiroshi Matsumoto, Satoshi Iwakami

## Abstract

Auxinic herbicides are critically important tools for weed control in modern agriculture. Although resistant weed populations have been increasing in recent years, the underlying mechanisms remain largely unknown. In a multiple-herbicide resistant line of the paddy weed *Echinochloa phyllopogon*, overexpression of catalytically promiscuous CYP81A P450 enzymes has been shown to confer resistance to diverse herbicides. Here, we investigated the role of these P450s in quinclorac resistance in *E. phyllopogon*. *Arabidopsis thaliana* plants expressing *CYP81A12* and *CYP81A21* genes, previously demonstrated to metabolize other herbicides, showed low but detectable levels of quinclorac resistance. A similar activity was also observed for CYP81A24, another promiscuous enzyme present in *E. phyllopogon*. Moreover, *Escherichia coli* strains expressing these genes produced putative hydroxylated metabolites of quinclorac. *In silico* analyses revealed that quinclorac binds stably near the catalytic heme center of CYP81A12, CYP81A21, and CYP81A24, but not CYP81A18, the importance of spatial alignment between the substrate recognition site and the heme center in determining metabolic activity and resistance potential. Consistent with the phenotype of resistant *E. phyllopogon*, *A. thaliana* expressing *CYP81A12* and *CYP81A21* produced significantly less ethylene following quinclorac treatment, linking P450-mediated metabolism to the suppression of quinclorac-induced phytotoxic responses. Finally, we demonstrated that CYP81A enzymes also confer resistance to other auxinic herbicides, florpyrauxifen-benzyl and 2,4-D. These findings provide new insights into the molecular basis of auxinic herbicide resistance and underscore the role of CYP81A P450s in metabolic detoxification and the prevention of herbicide-induced phytotoxic responses.

## INTRODUCTION

Weed control using herbicides has become an indispensable tool in modern crop production. However, continuous herbicide application imposes strong selection pressure, leading to the evolution of herbicide-resistant weed populations (Powles and Yu, 2010). Synthetic auxins, introduced as the first herbicide in 1945, have been widely used for decades and have remained effective with relatively low frequencies of resistance (Moreno-Serrano *et al*., 2024). Thus, while resistance to other herbicides has caused devastating problems, auxinic herbicides have continued to serve as valuable options for managing resistant weed populations. However, the recent deployment of genetically modified crops resistant to auxinic herbicides, along with the development of new selective auxinic herbicides for major crops, has resulted in a rapid increase in resistance reports (Todd *et al*., 2020). Addressing these resistant weeds requires a detailed understanding of their resistance mechanisms. Given the complex mode of action of auxinic herbicides, which mimic the plant hormone auxin, the molecular basis of resistance remains largely elusive. Unraveling these mechanisms is not only essential for sustainable weed management but also for advancing our understanding of auxin signaling and herbicide action.

As with other herbicide classes, resistance to auxinic herbicides can be classified into target-site resistance (TSR) and non-target-site resistance (NTSR) (Todd *et al*., 2020). While recent studies have begun to elucidate TSR mechanism for auxinic herbicides (LeClere *et al*., 2018; de Figueiredo *et al*., 2022; Montgomery *et al*., 2024), NTSR mechanisms remain poorly understood. Proposed mechanisms of NTSR include reduced translocation, enhanced metabolic detoxification (e.g., by cytochrome P450 monooxygenases), and the activation of detoxification enzymes such as β-cyanoalanine synthase (β-CAS) (Gaines *et al*., 2020). However, the specific molecular players involved in auxinic herbicide NTSR are still largely unknown, with only a few candidates such as CYP72A recently proposed (Wang *et al*., 2024).

Quinclorac, a widely used auxinic herbicide, strongly induces ethylene biosynthesis in plants, leading to the accumulation of hydrogen cyanide as a byproduct (Song *et al*., 2022). Because hydrogen cyanide is phytotoxic, its excessive accumulation following quinclorac treatment is considered a primary cause of plant death (Grossmann and Kwiatkowski, 2000). Accordingly, enhanced detoxification of hydrogen cyanide via β*-*CAS has been proposed as a resistance mechanism (Grossmann, 2010).

Previously, we investigated quinclorac resistance in a multiple-herbicide resistant (MHR) population of the paddy weed *Echinochloa phyllopogon* collected in California, USA in 1997. Interestingly, this population exhibited resistance to quinclorac despite no known history of exposure to auxinic herbicides (Yasuor *et al*., 2012). Our earlier studies showed that resistance to several non-auxinic herbicides in this population is conferred by a single genetic factor that drives the coordinated overexpression of multiple cytochrome P450 genes, i.e. *CYP81A12*, *CYP81A21*, and *CYP709C69* (Suda *et al*., 2023). In contrast, initial physiological analyses suggested that quinclorac resistance in this population could involve (1) increased β*-*CAS activity and (2) reduced ethylene production following herbicide treatment (Yasuor *et al*., 2012). Subsequent molecular analyses, however, ruled out the involvement of β*-*CAS (Chayapakdee *et al*., 2020), while resistance remained strongly associated with reduced ethylene production and resistance to herbicides with different modes of action. These findings raised the possibility that reduced ethylene production may result from rapid detoxification of quinclorac via overexpressed P450 enzymes.

In this study, we hypothesized that the catalytically promiscuous cytochrome P450s, CYP81A12 and CYP81A21, which are overexpressed in MHR *E. phyllopogon*, contribute to quinclorac resistance through metabolic detoxification. Using heterologous expression systems (*Arabidopsis thaliana* and*Escherichia coli*), *in vitro* metabolite analysis, and *in silico* structural modeling, we evaluated the quinclorac-metabolizing activity of these enzymes. Our results demonstrate that CYP81A overexpression leads to rapid quinclorac detoxification and suppression of ethylene production, providing a mechanistic explanation for auxinic herbicide resistance in this population. These findings contribute to a better understanding of NTSR mechanisms in weeds and offer new insights into how metabolic resistance can evolve to overcome diverse herbicide chemistries.

## RESULTS

### Quinclorac Response of *Arabidopsis thaliana* expressing *CYP81A12/21*

Quinclorac resistance of recombinant inbred lines (RILs) derived from a sensitive (S) and a MHR line of *E. phyllopogon* linked to the resistance to most of the herbicides, which is explained by co-regulation of multiple herbicide-metabolizing P450 (Chayapakdee *et al*., 2020), i.e. catalytically promiscuous CYP81A12 and CYP81A21, and highly substrate-specific CYP709C69. Although no activity to quinclorac was confirmed for CYP709C69 (Suda *et al*., 2023), CYP81A12 and CYP81A21 had not been analyzed yet. To confirm the role of these two P450s in quinclorac resistance, we tested the sensitivity of the previously established *Arabidopsis thaliana* lines expressing *CYP81A12* and *CYP81A21*. In our previous research, the lines exhibited marked resistance responses to many herbicides by 10 d cultivation on herbicide-supplemented MS media (Dimaano *et al*., 2020). However, in the case of quinclorac response, plant response was not very clear at 10 d (supplementary Fig. 1). Considering that *A. thaliana* is naturally tolerant to quinclorac, we further continued cultivation of the plants. Interestingly, the longer the plants were cultivated, the clearer the growth differences became. *A. thaliana* plants expressing these P450s grew better in quinclorac-containing medium compared to wild-type plants at 15 d (Fig. 1A).

**Figure 1.**
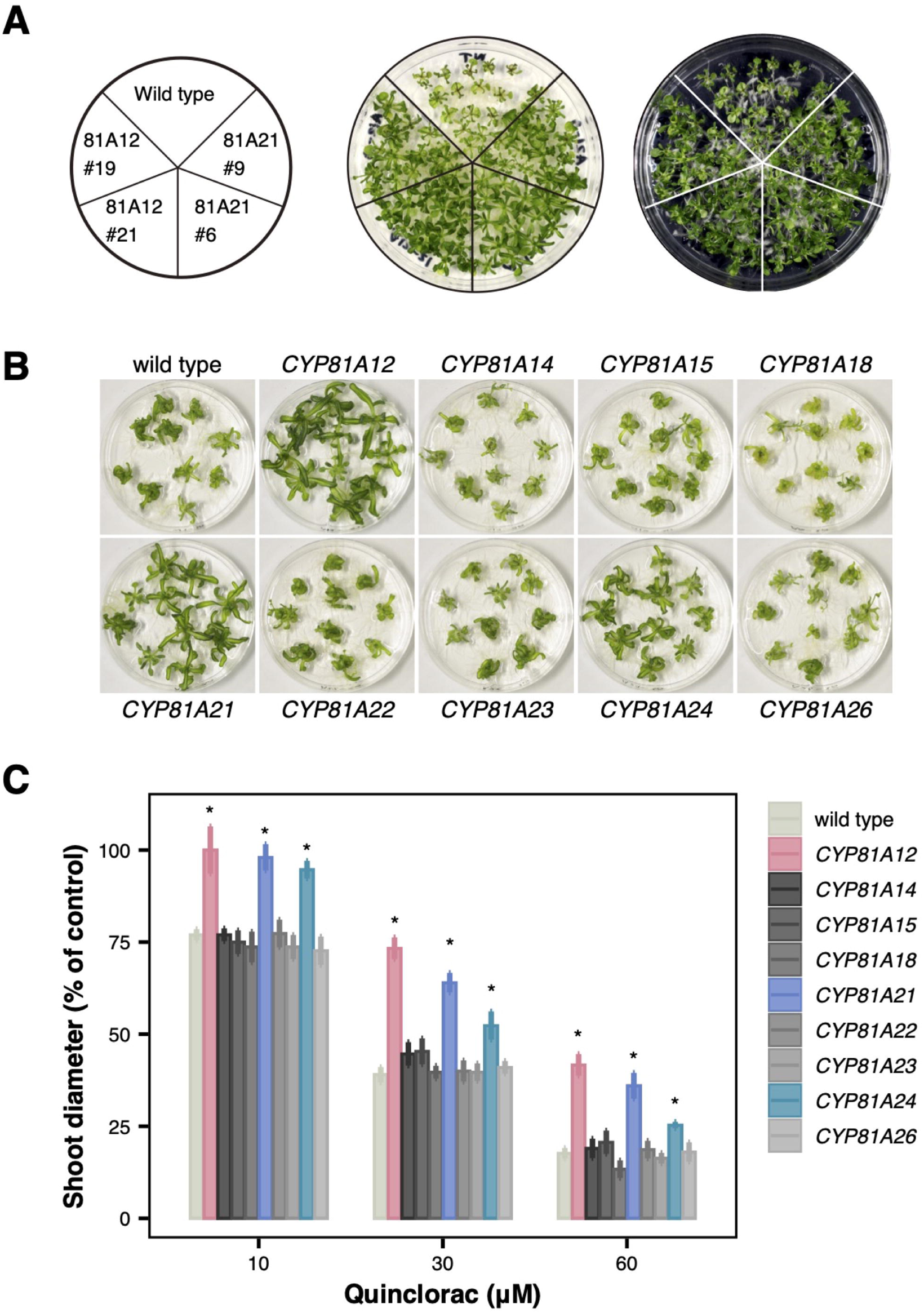
Quinclorac responses of *Arabidopsis thaliana* expressing CYP81A P450s of *Echinochloa phyllopogon.* (A) *A. thaliana* response to 100 μM quinclorac. Five-day-old plants were transferred to quinclorac medium and cultivated for additional 15 d. (B) *A. thaliana* response to 30 μM quinclorac. Plants were cultivated for 30 d. (B) Bar graph shows mean±SD of triplicates. * Significantly different from group wild type (*P*<0.05, Dunnett’s test).

Therefore, *A. thaliana* lines with higher expression, i.e., 81A12#21 and 81A21#6 (Iwakami *et al*., 2014), were further investigated under prolonged cultivation. The shoot diameters of the transformants were significantly larger under both 30 μM and 60 μM treatments at 30 d (Fig. 1B, C).

To further confirm the role of CYP81As in plant response to quinclorac, other *CYP81A* genes present in *E. phyllopogon*, i.e., *CYP81A14*/*15*/*18*/*22*/*23*/*24*/*26*, which are not associated with resistance in the MHR line, were also investigated. The results revealed that, in addition to *CYP81A12* and *CYP81A21*, *CYP81A24* also conferred mild resistance to quinclorac in *A. thaliana* (Fig. 1B, C). These findings indicate that overexpression of CYP81A P450 in plants can confer resistance to quinclorac.

### Role of CYP81As in quinclorac metabolism

To determine whether CYP81As metabolize quinclorac, we performed an *E. coli* whole-cell metabolism assay using a previously established protocol. Cultures were supplemented with 0.2 mM quinclorac and incubated for 24 hours, followed by LC-MS/MS analysis of the culture medium. We tested three CYP81A enzymes that conferred quinclorac resistance in *A. thaliana* and two others, CYP81A15 and CYP81A18, which did not confer resistance. A peak corresponding to hydroxylated quinclorac was monitored in multiple reaction monitoring (MRM) mode. A clear peak with a retention time of 5.1 min was identified in the culture solution of *E. coli* with *CYP81A12*, *CYP81A21*, and *CYP81A24* (Fig. 2), in accordance with the results in *A. thaliana*. The findings indicate that CYP81A enzymes have hydroxylation activity toward quinclorac.

**Figure 2.**
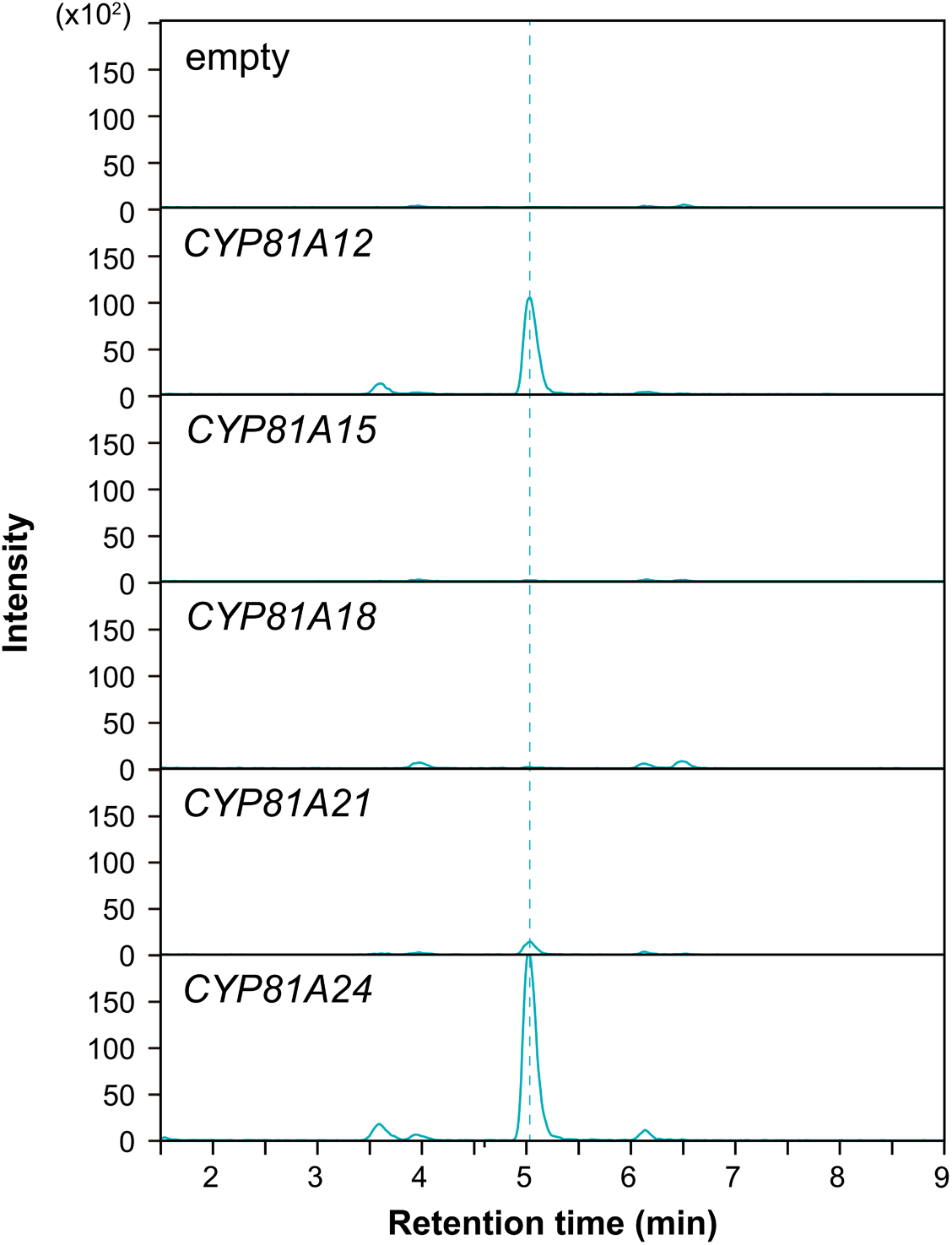
Detection of putative hydroxylated quinclorac in *CYP81A* expressing *E scherichia coli*.

### Quinclorac/CYP81As interactions via *in silico* study

To investigate the quinclorac-metabolizing activity of CYP81As in *E. phyllopogon*, we conducted an *in silico* analysis of their interactions with quinclorac. Due to the absence of available crystal structures, the 3D structures of the CYP81As were generated using the Robetta server. The geometry and stereochemistry of these models were validated using Ramachandran plots (Fig. S2). The structural models revealed the presence of six putative substrate recognition sites (SRSs), highlighting the high degree of similarity between CYP81A12, CYP81A21, and CYP81A24, with approximately 75% sequence identity. In contrast, CYP81A18, which does not confer quinclorac resistance, shares only 50% sequence identity (Figs.S3, S4). We first employed molecular docking and all-atom molecular dynamics (MD) simulations to investigate further the molecular basis of the metabolism, where hydrogen atoms are omitted by being included in groups such as CH_3_. The molecular docking results demonstrated that quinclorac could bind to the active sites of the minimized structures of CYP81As located near the heme catalytic region. Consistent with the *A. thaliana* and *E. coli* studies, the binding energy scores for quinclorac interacting with CYP81A12 (−7.6 kcal/mol), CYP81A21 (−7.3 kcal/mol), and CYP81A24 (−7.9 kcal/mol) were more favorable compared with CYP81A18 (−6.6 kcal/mol), which is the negative control in this study (Fig. S5). These interactions are influenced by the hydrophobicity, hydrophilicity, and amphipathicity of the SRSs, which determine how quinclorac accesses to and is stabilized in the binding pocket. Several residues within the SRS domains interacted with the core structure of quinclorac through van der Waals (vdW), π-π, and π-alkyl interactions, consistent with previous studies in other P450s (El-Awaad *et al*., 2016; Watanabe *et al*., 2017). In CYP81A12 and CYP81A21, polar residues facilitated interactions with the carboxyl group of quinclorac through hydrogen bonds or ionic interactions, guiding it toward the heme. CYP81A24, with the highest amphipathicity (8.00%), exhibited a balance of hydrophobic and hydrophilic residues, allowing dynamic binding and stabilizing both aromatic and polar moieties in quinclorac. In contrast, CYP81A18, with the highest hydrophobicity (54.67%), interacted primarily with quinclorac through vdW forces, which likely restricted its movement deeper into the pocket and hindered access to the heme, potentially affecting catalytic activity.

All-atom MD simulations were conducted for the quinclorac/CYP81A complexes to assess the binding stability and conformational dynamics over 300-ns trajectories (Video 1). The root-mean-square deviation (RMSD) analysis of quinclorac and the backbone of CYP81As indicated high stability throughout the simulations (Fig. 3A). Whereas, in the CYP81A21 and CYP81A24 systems, quinclorac exhibited slight fluctuations due to the rotational freedom of its carboxyl group. Analysis of the distance between the center of mass of quinclorac and the heme iron (*d*_Fe-Quinclorac_) throughout the MD trajectory suggested that quinclorac gradually moved closer to the catalytic site of the heme in CYP81A12 (∼5 Å), CYP81A21 (∼7 Å), and CYP81A24 (∼6 Å), whereas it remained more distant in CYP81A18 (∼11 Å) (Fig. S5B; Video 1). The hydration dynamics around heme in the pocket were analyzed across four systems by evaluating the total number of water molecules (#Water) within the inner (0–3.4 Å) and outer (3.4–5 Å) hydration shells of Fe-heme. CYP81A12 and CYP81A21 showed low water occupancy at the Fe-heme, indicating a less solvent-exposed or more hydrophobic environment. CYP81A24 exhibited moderate water occupancy with slight fluctuations, suggesting periodic changes in solvent accessibility. Due to the motion of quinclorac, we found that water can access the pocket of CYP81A18 and move close to the Fe-heme, which is higher than the other systems, leading to the loose binding of quinclorac in the binding pocket (Fig. S5C). The CYP81A18 system revealed a unique two-dimensional free energy landscape (2D-FEL) pattern characterized by two global minima across a broad chemical space. These minima were observed at both lower and higher water densities within the binding pocket compared to the other systems (Fig. 3B). In contrast, the other three systems displayed a narrower 2D-FEL with a single global minimum, suggesting higher ligand stability. During the MD simulation, CYP81A12 exhibited the largest binding pocket volume (average from the last 50 ns: 130.56 ± 9.54 nm^3^) and the highest range of structural fluctuations, supporting its superior metabolic activity (Fig. 3C and S5). The larger, more flexible pocket likely enhances substrate binding and positioning, improving catalytic efficiency for quinclorac metabolism. On the other hand, CYP81A18 had a smaller, narrower binding pocket (average from the last 50 ns: 48.82 ± 21.17 nm^3^), which likely restricted substrate access to the heme and hindered proper positioning, resulting in reduced activity.

**Figure 3.**
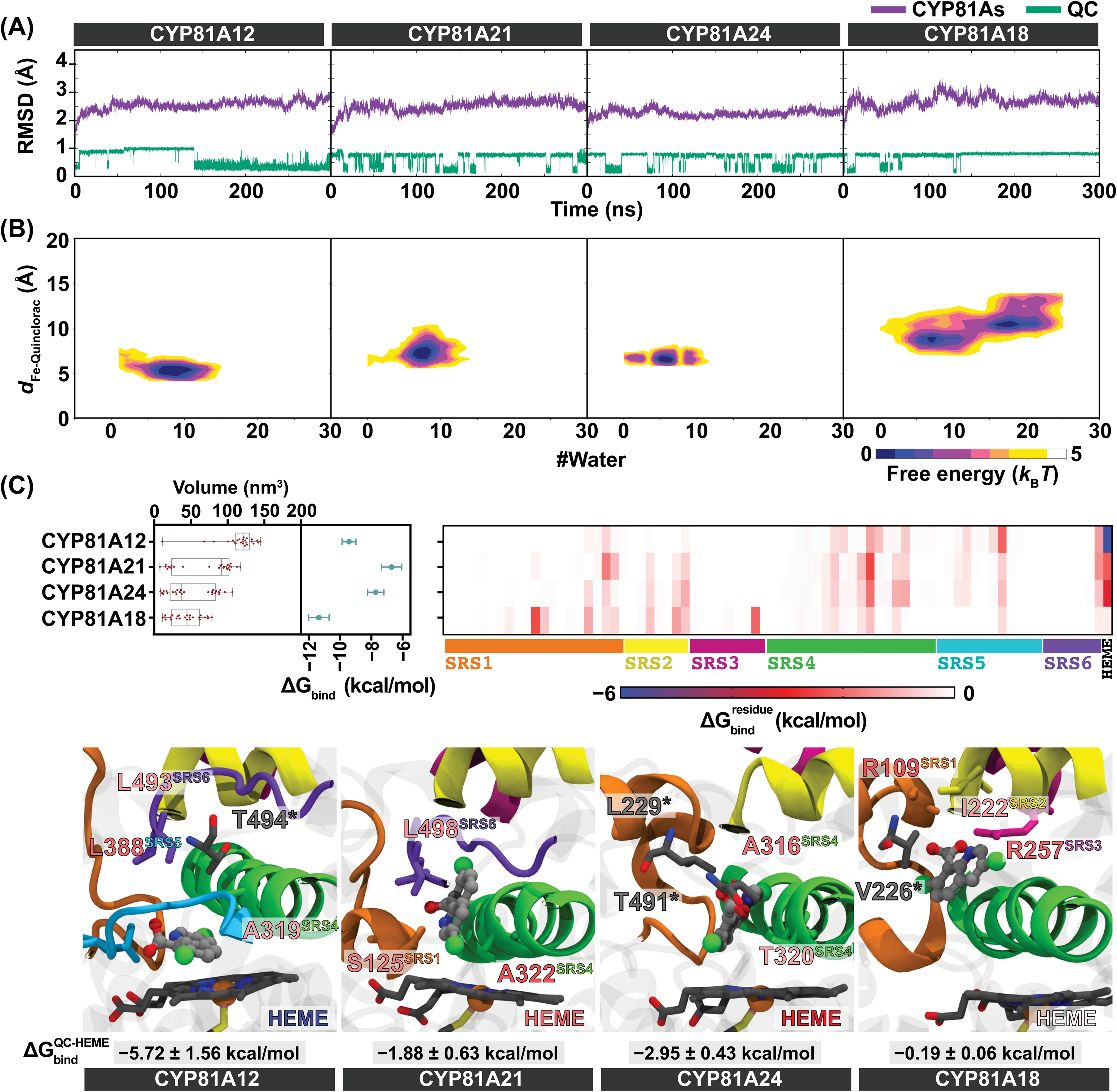
Structural dynamics and binding profiles of quinclorac in CYP81A12, CYP81A21, CYP81A24, and CYP81A18 variants. (A) Cα-RMSD and quinclorac–heme distance (*d*_Fe–Quinclorac_) over 300 ns MD simulations. (B) Two-dimensional free energy landscapes (2D-FEL) showing *d*_Fe–Quinclorac_ versus the number of water molecules in the binding pocket (#Water). (C) Binding pocket volume, binding free energy (ΔG ), and per-residue energy decomposition (ΔG^residue^_bind_) calculated by the MM/GBSA method, along with representative binding poses.

The binding (ΔG _bind_) and decomposition free energy 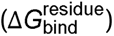 calculations, derived from the final 50 ns of MD trajectory, reveal that quinclorac binds more effectively to the SRSs of CYP81A18 (−11.35 ± 0.65 kcal/mol) than CYP81A12 (−9.42 ± 0.44 kcal/mol), CYP81A21 (−6.70± 0.64 kcal/mol), and CYP81A24 (−7.72 ± 0.52 kcal/mol) (Table S2). However, in CYP81A18, quinclorac is located far from the catalytic center (Fig. 3C). Consistent with the interaction profile from 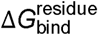 quinclorac in CYP81A18, it moves away from the heme and interacts stably with R257 on SRS3. In CYP81A21 and CYP81A24, quinclorac forms stable π-alkyl interactions with residues A322 and A316 in SRS4 (Fig. 3C). Despite fewer contributing residues in CYP81A12, it exhibits the strongest 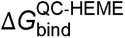 (−5.72 ± 1.56 kcal/mol), indicating a potential for high quinclorac-metabolizing activity. Based on the interaction profile (Fig. 3C), quinclorac consistently interacts with the α-helix in the SRS4 and leucine in SRS6 in all systems. The close proximity of these interactions to the heme group in CYP81A12, CYP81A21, and CYP81A24 indicates stronger binding affinities and greater potential for quinclorac metabolism. This could explain why these CPY81As are more efficient at quinclorac metabolism, as their interaction networks are better aligned with the catalytic core. The interaction shift observed in CYP81A18, involving regions distant from the heme, may underlie its reduced capacity for quinclorac metabolism, reinforcing the specificity of certain CYP81A proteins in quinclorac resistance. These findings suggest that the spatial arrangement of SRSs in relation to the heme group is critical for the efficiency of herbicide metabolism, offering valuable insights into the molecular mechanisms underlying resistance.

### Association of quinclorac resistance and ethylene production

In our previous study, the complete association between quinclorac resistance and reduced ethylene production was observed in the RILs of *E. phyllopogon* (Chayapakdee *et al*., 2020). Together with the above findings that CYP81A12 and CYP81A21 metabolize quinclorac, we hypothesized that reduced ethylene biosynthesis in resistant *E. phyllopogon* is caused by the decreased quinclorac level due to faster quinclorac metabolism. To test this hypothesis, ethylene production in *A. thaliana* lines expressing *CYP81A12*/*21* was measured after quinclorac application. *Arabidopsis thaliana* plants grown on MS medium for 30 d were transferred to quinclorac solutions (0, 300, and 1000 µM) for 12 h, and then kept in glass vials (Fig. 4A). After 96 h, ethylene levels were measured using gas chromatography. The results demonstrated that *A. thaliana* carrying *CYP81A12*/*21* produced significantly less ethylene (578 and 892 nl gFW^-1^, respectively) than wild-type plants (2,791 nl gFW^-1^) (Fig. 4B). This study confirmed that upregulation of *CYP81A12*/*21* enhances quinclorac degradation and reduces ethylene production in plants.

**Figure 4.**
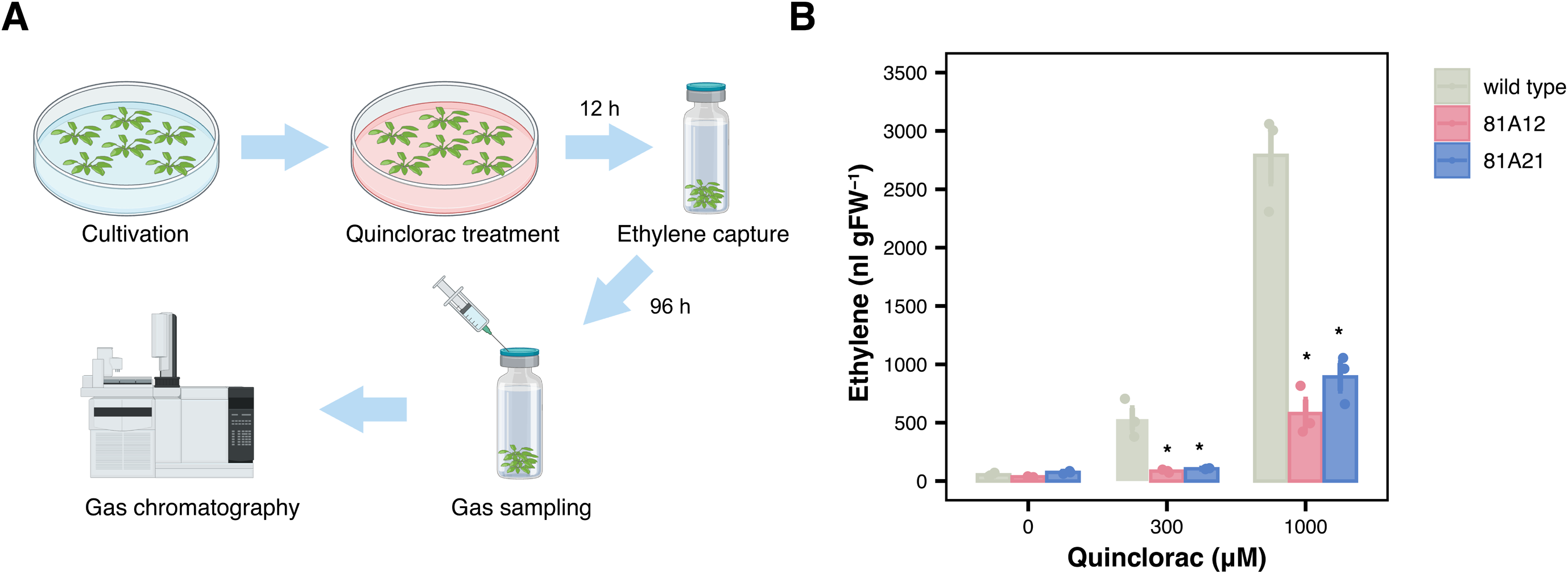
Ethylene production after quinclorac treatment in *Arabidopsis thaliana*. (A) Methodology of ethylene quantification. (B) Ethylene levels in quinclorac treated plants. Data showed mean±SD of triplicates (3 vials, 3 plants/vial). * Significantly different from group wild type (*P*<0.05, Dunnett’s test).

### Cross-resistance to auxin herbicides by CYP81A P450s

Having identified the metabolic function of CYP81A toward quinclorac, we next assessed its activity against other auxinic herbicides. Florpyrauxifen-benzyl and 2,4-D were selected as representative auxinic herbicides for testing, as 2,4-D is one of the earliest and still widely used auxinic herbicides, and florpyrauxifen-benzyl represents a new generation of auxin herbicides that, unlike quinclorac, is the first to be applicable for use in rice cultivation (Fig. 5A). Florpyrauxifen-benzyl and 2,4-D belong to the 6-arylpicolinate and phenoxycarboxylate classes, respectively, which are structurally distinct from quinclorac, a quinolinecarboxylate (Moreno-Serrano *et al*., 2024). *Arabidopsis thaliana* lines expressing the quinclorac-metabolizing genes *CYP81A12* and *CYP81A21* exhibited resistance to both florpyrauxifen-benzyl and 2,4-D (Fig. 5B, 6C).

**Figure 5.**
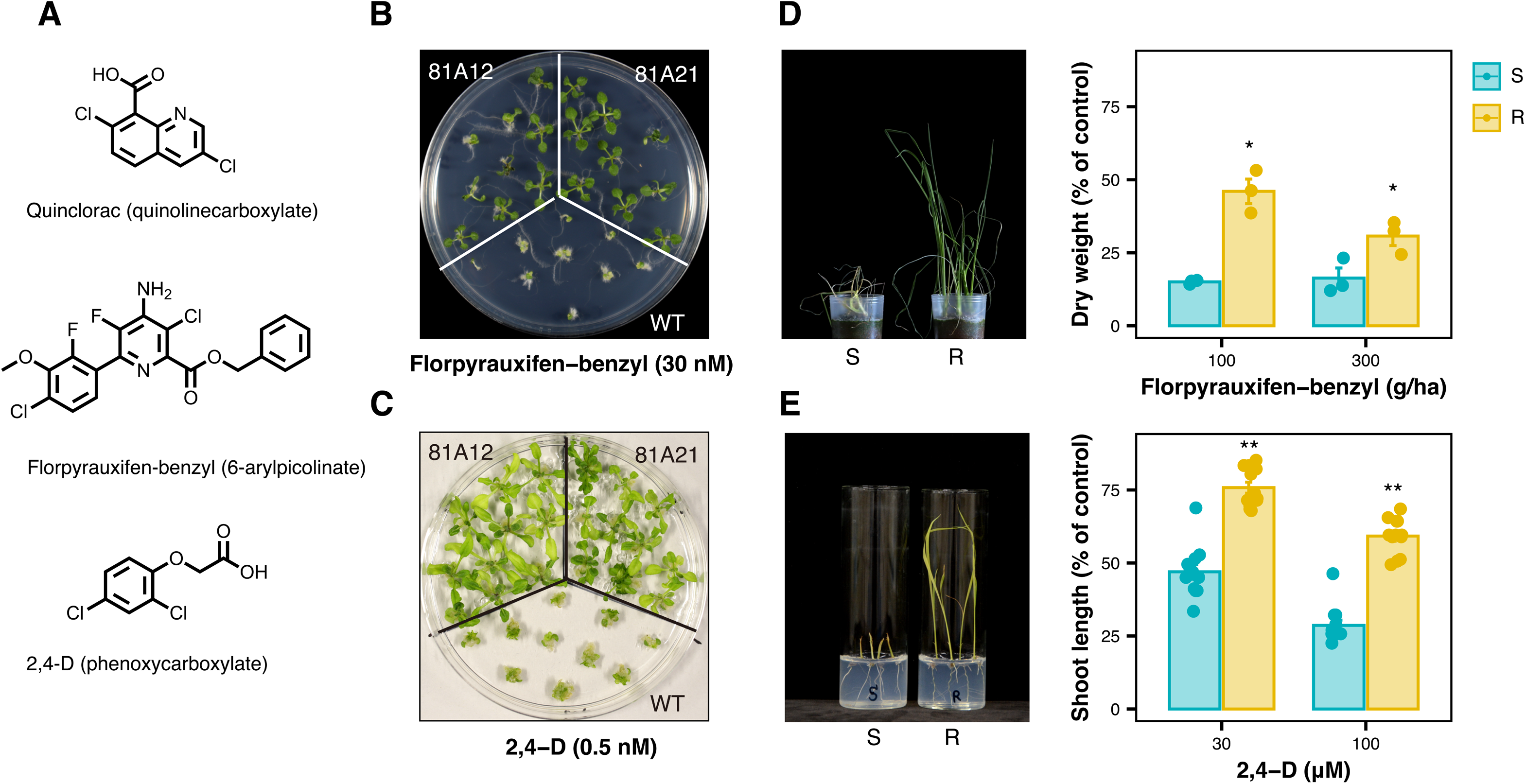
Role of CYP81A P450s in plant sensitivities to other auxinic herbicides. (A) Chemical structure of florpyrauxifen-benzyl and 2,4-D. (B) Florpyrauxifen-benzyl responses of *Arabidopsis thaliana* expressing *CYP81A12* and *CYP81A21*. Fourteen-day-old plants. (C) 2,4-D responses of *A. thaliana* expressing *CYP81A12* and *CYP81A21*. Thirty-day-old plants. (D) Florpyrauxifen-benzyl responses of sensitive and multiple-herbicide-resistant *Echinochloa phyllopogon*. Bars, ± SE (n = 3). * significant difference of *P*<0.05 (Welch’s *t*-test). (E) 2,4-D responses of sensitive and multiple-herbicide-resistant *E. phyllopogon*. Bars, ± SE (n = 3). ** significant difference of *P*<0.01 (Welch’s *t*-test).

Based on these results, we hypothesized that the MHR *E. phyllopogon* overexpressing these genes would exhibit resistance to these herbicides. The MHR line showed clear resistance to florpyrauxifen-benzyl when treated at the standard application rate (300 g/ha). Although 2,4-D is generally less effective against grass species, high concentrations can still inhibit their growth. The MHR line also exhibited marked resistance to 2,4-D, similar to its response to florpyrauxifen-benzyl. These results indicate that CYP81A P450s are capable of detoxifying auxinic herbicides with diverse chemical structures. This is consistent with their previously reported ability to metabolize herbicides with differing modes of action.

## Discussion

In recent years, resistance to auxinic herbicides has emerged in several weed species. However, aside from a few cases involving target-site resistance (TSR), the underlying molecular mechanisms remain poorly understood. Among these herbicides, quinclorac has long been considered metabolically stable in plants, and its resistance mechanisms have not been thoroughly investigated. In this study, we focused on elucidating the basis of quinclorac resistance in the MHR *E. phyllopogon* population. A previous study has proposed mechanisms such as reduced ethylene production and the activation of β-CAS as contributing factors (Yasuor *et al*., 2012). However, our prior work demonstrated that β-CAS overexpression in *E. phyllopogon* is not associated with quinclorac resistance (Chayapakdee *et al*., 2020). Here, we show that CYP81A-mediated quinclorac metabolism underlies the reduction in ethylene production, as evidenced by experiments using heterologous expression systems in *A. thaliana* and *E. coli*. The low resistance level conferred by *CYP81A12* and *CYP81A21* in *A. thaliana* is consistent with the relatively low quinclorac resistance observed in the MHR *E. phyllopogon* population. Taken together, these findings provide strong evidence that cytochrome P450-mediated metabolism plays a key role in quinclorac resistance in *E. phyllopogon*.

The role of catalytically promiscuous P450s in metabolic cross-resistance was first demonstrated in *E. phyllopogon* (Iwakami *et al*., 2014; Iwakami *et al*., 2019; Guo *et al*., 2019). Later, the detoxification capabilities of *E. phyllopogon* CYP81As have been extensively evaluated, revealing that CYP81A12, CYP81A21, and CYP81A24 can metabolize a wide range of herbicides used in modern agriculture (Dimaano *et al*., 2020). This study further revealed that the promiscuity of CYP81A P450s extends to auxinic herbicides. Notably, florpyrauxifen-benzyl, a 6-arylpicolinate synthetic auxin, was first commercialized in 2018—nearly two decades after the collection of the MHR *E. phyllopogon* population. The observation that this population already exhibits reduced sensitivity suggests that broad-spectrum metabolic resistance mechanisms can predate the deployment of new herbicide chemistries, posing a serious challenge to proactive resistance management. CYP81A24 had already been shown to metabolize herbicides targeting six distinct modes of action—acetolactate synthase, phytoene desaturase, 4-hydroxyphenylpyruvate dioxygenase, 1-deoxy-D-xylulose 5-phosphate synthase, acetyl-CoA carboxylase, and protoporphyrinogen oxidase (Dimaano and Iwakami, 2021; Iwakami *et al*., 2019). It is now evident that CYP81A24 also metabolizes herbicides targeting a seventh mode of action, synthetic auxin.

Meanwhile, the underlying reasons why these P450s can detoxify such a diverse array of herbicides remained unclear. In this study, *in silico* analyses were conducted to investigate the metabolic mechanism of quinclorac. By comparing CYP81A enzymes capable of metabolizing quinclorac (CYP81A12/21/24) with those unable to do so (CYP81A18), we explored the structural and functional characteristics of CYP81A enzymes. The docking studies and molecular dynamics simulations corroborated the experimental findings, showing that CYP81A12, CYP81A21, and CYP81A24 have stronger binding affinities for quinclorac than CYP81A18. The water molecules near Fe in CYP81A18 (Fig.S5) may obstruct substrate access and reduce the conformational flexibility of quinclorac, limiting its catalytic efficiency compared to the others. Therefore, the solvent environment could be crucial for the activity of CYP81A (Fig. 3C, S5C, and S6). The pocket volume analysis showed that quinclorac bound to CYP81A12 has the largest pocket, allowing greater flexibility and easier access to the heme center. In contrast, CYP81A18 has the smallest pocket and strongest binding to surrounding residues, restricting quinclorac’s movement toward the heme and reducing metabolic activity due to interactions with SRS1, SRS2, and SRS4 (Fig. 3C). The larger pocket and lower water content in CYP81A12, CYP81A21, and CYP81A24 likely enhance its metabolic performance. The detailed analysis of SRSs and the binding free energy profiles sheds light on the molecular mechanisms that underlie quinclorac resistance, particularly the key residues that facilitate effective herbicide metabolism. However, the mode of quinclorac metabolism by CYP81As, e.g., the specific hydroxylation site, remains unclear. Several high computational approaches, i.e., quantum mechanics/molecular mechanics (QM/MM) calculation or cluster model analysis, can be further utilized to confirm the reaction of CYP81As-quinclorac at the quantum level. Furthermore, it is essential to evaluate the detoxification mechanisms for other herbicides. While plant P450 enzymes were long thought to exhibit high substrate specificity, recent discoveries have identified P450s, including CYP81A and others, with broad substrate versatility (Werck-Reichhart, 2023). Computational simulations hold promise for uncovering the structural and functional characteristics that enable these P450s to process diverse substrates.

One of the significant insights revealed through a series of studies on MHR *E. phyllopogon* is its contribution to understanding the mode of action of synthetic auxins. While the reception and signaling pathways of endogenous auxins are well understood, the mechanisms by which synthetic auxins exert their effects remain unclear (Gaines, 2020). It is widely believed that ethylene production induced by synthetic auxin application plays a central role in the processes leading to plant death. Our analysis of *A. thaliana* expressing the quinclorac-metabolizing P450 revealed a negative correlation between ethylene release and synthetic auxin perception, which supports earlier studies highlighting the link between ethylene evolution and plant death (Grossmann, 2010). However, our research strongly suggests that hydrogen cyanide, a by-product of ethylene biosynthesis previously considered a key factor in plant death, is not involved in this process (Chayapakdee *et al*., 2020). Recent studies have also suggested the possibility that the accumulation of abscisic acid (ABA), independent of ethylene evolution, may contribute to plant death (McCauley *et al*., 2020). Further investigation of ABA-related responses using our *E. phyllopogon* materials may provide additional insights into the mode of action of synthetic auxins.

Our research has progressively uncovered key herbicide-detoxifying genes involved in MHR in *E. phyllopogon*. In particular, we have identified the overexpression of promiscuous cytochrome P450 enzymes, as well as the coordinated upregulation of multiple detoxifying enzymes. These findings have significantly advanced our ability to predict cross-resistance patterns associated with metabolism-based herbicide resistance. However, despite these advances, the regulatory mechanisms driving the simultaneous overexpression of these detoxification genes remain elusive. The strikingly similar expression patterns observed across independently evolved MHR populations suggest the involvement of common upstream regulatory factors (Suda *et al*., 2023). With the growing availability of genomic resources, including multiple genome assemblies (Wu *et al*., 2022; Sato *et al*., 2023) and recombinant inbred lines (RILs) (Iwakami *et al*., 2014), *E. phyllopogon* is emerging as a powerful model for studying MHR in weeds. Harnessing these resources will be essential for achieving a comprehensive understanding of metabolism-based herbicide resistance—a critical mechanism in modern weed management—and for informing the development of effective and sustainable control strategies.

## MATERIALS AND METHODS

### Plant material

Two inbred lines of *E. phyllopogon* were used: MHR line (line 511) and S line (line 401). These MHR and S lines were originally collected in 1997 from rice fields in California’s Sacramento Valley and were self-pollinated for three consecutive generations (Tsuji *et al*., 2003). The MHR line exhibits resistance to acetolactate synthase inhibitors (Yasuor *et al*., 2009), acetyl-CoA carboxylase inhibitors (Iwakami *et al*., 2019), a DXS inhibitor (Yasuor *et al*., 2008), very-long-chain fatty acid elongase inhibitors (Fischer *et al*., 2000), in addition to auxinic herbicides shown in this study.

Transgenic *A. thaliana* plants overexpressing each gene from the *CYP81A* subfamily were generated previously (Iwakami *et al*., 2014; Guo *et al*., 2019). Herbicide sensitivity of each line was described previously (Dimaano *et al*., 2020). 81A12#21 and 81A21#6 were used unless stated otherwise.

### Herbicide sensitivity assays

Unless otherwise stated, herbicide sensitivity of *A. thaliana* was evaluated as described previously (Guo *et al*., 2019). Briefly, sterile seeds were transferred to Murashige and Skoog (MS) medium plates containing herbicides and kept in a growth chamber at 22°C under constant light (20 - 40 µmol m^−2^s^−1^) for up to 30 d. For the experiment shown in Fig. 1A and S1, 5-day-old seedlings germinated on quinclorac-free MS medium were transferred to MS medium containing quinclorac.

Herbicide sensitivity assays for the S and MHR lines of *E. phyllopogon* were conducted using solid medium culture. Seeds from the S and MHR lines were sterilized with 2.5% (w/v) sodium hypochlorite for 15 min, further sterilized with 0.2% (w/v) sodium hypochlorite for 30 min and washed three times with sterile water. The seeds were germinated on wet filter paper in petri dishes for three d in a growth chamber set at 25°C with a 12-hour photoperiod (250 – 300 µmolm^−2^ s^−1^). The seedlings were then transferred to MS solid medium containing herbicides.

For foliar spray treatment of florpyrauxifen-benzyl, germinated *E. phyllopogon* seeds were transplanted to soil with four plants per pot and cultivated under LED with 13-h photoperiod and 27±°C. Plants under 3.8 to 4-leaf-stage were sprayed according to Iwakami et al. (2024). Florpyrauxifen-benzyl (Loyant 2.7% EC; Corteva Japan Ltd) were treated at the 1/3X and 1X of the recommended rate (50 g a.i./ha). Plants were collected at 16 d after application and dried for 3 d at 80 °C for dry weight measurement.

### Quantification of ethylene production

Ethylene production was determined according to the method described previously (Chayapakdee *et al*., 2020) with some modifications. *Arabidopsis thaliana* plants were germinated in solid MS medium under the same conditions as above for 30 d. Then, the roots of grown plants were immersed in quinclorac solution for 12 h and three plants were transferred to a 30 mL glass vial containing filter paper and 2 mL distilled water. The vial was sealed tightly and maintained under germination conditions. After 96 h incubation, 1 mL gas samples were withdrawn by syringe from the vial and quantified for ethylene by gas chromatography (GC-17A; Shimadzu Corp., Kyoto, Japan). The GC contained a glass column packed with Unipak S (GL Sciences Inc., Tokyo, Japan) and a hydrogen flame ionization detector. The temperatures of the chromatograph oven, injection, and detector were 60, 120, and 120°C, respectively. Nitrogen carrier gas, hydrogen gas, and air pressures were adjusted to 50 mL min^−1^, 60 kPa, and 50 kPa, respectively. The quinclorac concentrations applied to the plants were 0, 300, or 1000 μM. Ethylene production was measured in triplicate.

### Quinclorac metabolism

Details of *E. coli* expressing each *CYP81A* gene from *E. phyllopogon* was described in our previous study (Dimaano *et al*., 2020). Briefly, the *E. coli* C41(DE3) lines carry pET28 vector with a *CYP81A* gene, pCDF vector with *ATR2* and 5-aminolevurinic acid synthase (*HemA*) genes, and pGro7 with chaperonin gene. *E. coli* colonies were inoculated into LB containing glucose (1% w/v), kanamycin (50 µg/ml), streptomycin (50 µg/ml), and chloramphenicol (34 µg/ml), and cultured at 37°C overnight. A 10 µl aliquot was transferred to 1 ml of auto-induction medium, followed by culture at 26°C for 24 hours. Quinclorac (0.2 mM) and arabinose (2 mg/ml) were added, and the culture was continued at 30°C for 24 hours.

Samples were analyzed using liquid chromatography-tandem mass spectrometry (LC-MS/MS) with a COSMOSIL 2.5C18-MS-II column under the following conditions: mobile phase A, 0.1% formic acid in water; mobile phase B, acetonitrile; and a 10–60% linear gradient of B for 8 min and 60% B for 1 minute at a flow rate of 0.4 ml/min. Mass spectrometry was conducted in positive or negative ion mode using an LCMS-8030 (Shimadzu, Japan), with MRM used to detect hydroxylated quinclorac metabolite under the following conditions: positive-ion mode with MRM transition of *m*/*z* 258 [M+H]+ to *m*/*z* 177.

### In silico analyses

#### CYP81As Protein Structure Modeling

The protein sequences of CYP81As were derived from previously identified nucleotide sequences (Iwakami *et al*., 2014). The 3D structural models were generated using integrated template-based and *de novo* prediction techniques via the Robetta server (Kim *et al*., 2004). Hydrogen atoms were then added to the protein models, and heme groups were positioned within the active sites through structural superimposition with a protein template (PDB code: 6VBY) (Zhang *et al*., 2020). Structural minimization of all models was performed using the ff19SB AMBER force field (Maier *et al*., 2015) for standard amino acids, while the penta-coordinate ferric high-spin (heme) was minimized using parameters from previous research implemented in AmberTools 21 (Shahrokh *et al*., 2012).

#### Molecular Docking

To predict the possible conformation and binding affinity between quinclorac and CYP81As, the minimized structures of CYP81As served as templates for molecular docking using Autodock Vina 1.1.2 (Trott and Olson, 2010). The structure of quinclorac was constructed using GaussView 6.0 and optimized with Gaussian 16 using the DFT B3LYP/6-31G* basis set (Frisch *et al*., 2016). Ligand and protein structures were charged and converted to PDBQT format using AutoDockTools (Morris *et al*., 2009). The binding site in CYP81As was defined around the heme, surrounded by SRS1 to SRS6, with grid dimensions of 15 x 15 x 15 Å, defined by the AutoGrid 4.0.0 module (Morris *et al*., 2009). The ligand-binding pose and interactions were analyzed in 2D and 3D using Discovery Studio Visualizer Software (BIOVIA, 2021) and Chimera USCF (Pettersen *et al*., 2004). The quinclorac/CYP81As complexes with the lowest binding energies were selected for MD simulations to assess the stability of ligand binding and explore molecular mechanisms.

#### Molecular Dynamics Simulation of Quinclorac/CYP81As Complexes

The initial structures of the quinclorac/CYP81As complexes were subjected to MD simulations using the AMBER20 software package (Case *et al*., 2020). Each complex was first prepared by examining its protonation state using the PDB2PQR web tools (Dolinsky *et al*., 2004). Hydrogen atoms and ions were then added to each system, and hydrogen atoms were minimized using the SANDER tool from AmberTools 21 (Case *et al*., 2020). Each complex was solvated with a TIP3P explicit water model, maintaining a cutoff distance of 12 Å from the protein’s surface. This was followed by 3,000 steps of steepest descent and 3,000 steps of conjugate gradient minimization.

The systems were gradually heated to 300 K over 20 ps under periodic boundary conditions with a canonical ensemble (NVT). The simulations were then continued at 300 K for 300 ns under an isothermal-isobaric ensemble (NPT). The stability of the systems was examined by monitoring the root-mean-square deviation (RMSD) and the displacement between iron in the heme and quinclorac (*d*_Fe-Quinclorac_) over the 300 ns simulation calculated by the CPPTRAJ module implemented in AmberTools 21. The binding pocket volume of CYP81A enzymes was calculated using POVME3.0(1) with default parameters. The grid was centered 2 Å above the heme group, which corresponds to the enzymatic active site. A spherical radius of 15 Å was used to define the pocket region. For each enzyme, 30 representative structures were extracted from the 300-ns MD trajectory in PDB format at 10-ns intervals. These structures were used to capture the dynamic changes in pocket volume during the simulation. The two-dimensional free energy landscape (2D-FEL) profile was performed based on the Markov state models (MSMs) using PyEMMA. From the last 50 ns of each system, a total of 5,000 snapshots were extracted and used for per-residue decomposition free energy calculations using the molecular mechanics generalized Born solvent-accessible surface area (MM/GBSA) method (Kollman *et al*., 2000), which identifies key amino acids involved in ligand binding to the protein.

## AUTHOR CONTRIBUTIONS

SI, PC and KH designed the research; PC, ND, and SI performed plant experiments; TY performed *E. coli* and LC-MS/MS analysis; KH performed *in silico* analysis; PC, KH, and SI wrote the paper; and ND, Y Shigeta, Y Sunohara, and HM critically revised this manuscript. All authors read and approved the manuscript.

## ACKNOWLEDGMENTS

This work was supported in part by JSPS KAKENHI (Grant Numbers: JP15H06072, JP17K15234, JP19H02955, JP23K23612 to SI), JST PRESTO (Grant Number JPMJPR24N2 to SI), and by partial support from Chiang Mai University. We are grateful to BASF for providing quinclorac.

---

**Video 1.** The MD trajectory of quinclorac binding in CYP81A variants over 300 ns shows together with the quinclorac–heme distance (*d*_Fe-Quinclorac_).

