## Supplementary material for "Promiscuous cytochrome P450s confer metabolic resistance to synthetic auxin herbicides in the weedy grass *Echinochloa phyllopogon*": Fig. S1

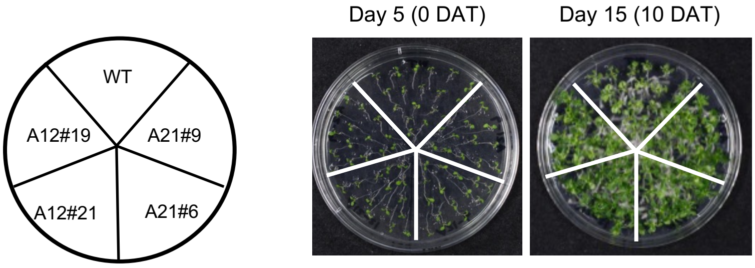


**Figure S1. Quinclorac responses of *Arabidopsis thaliana* expressing *CYP81A12* and *CYP81A21* of *Echinochloa phyllopogon.***

Five-day-old plants were transferred to 100 μM quinclorac medium.
