## Supplementary material for "Promiscuous cytochrome P450s confer metabolic resistance to synthetic auxin herbicides in the weedy grass *Echinochloa phyllopogon*": Fig. S2

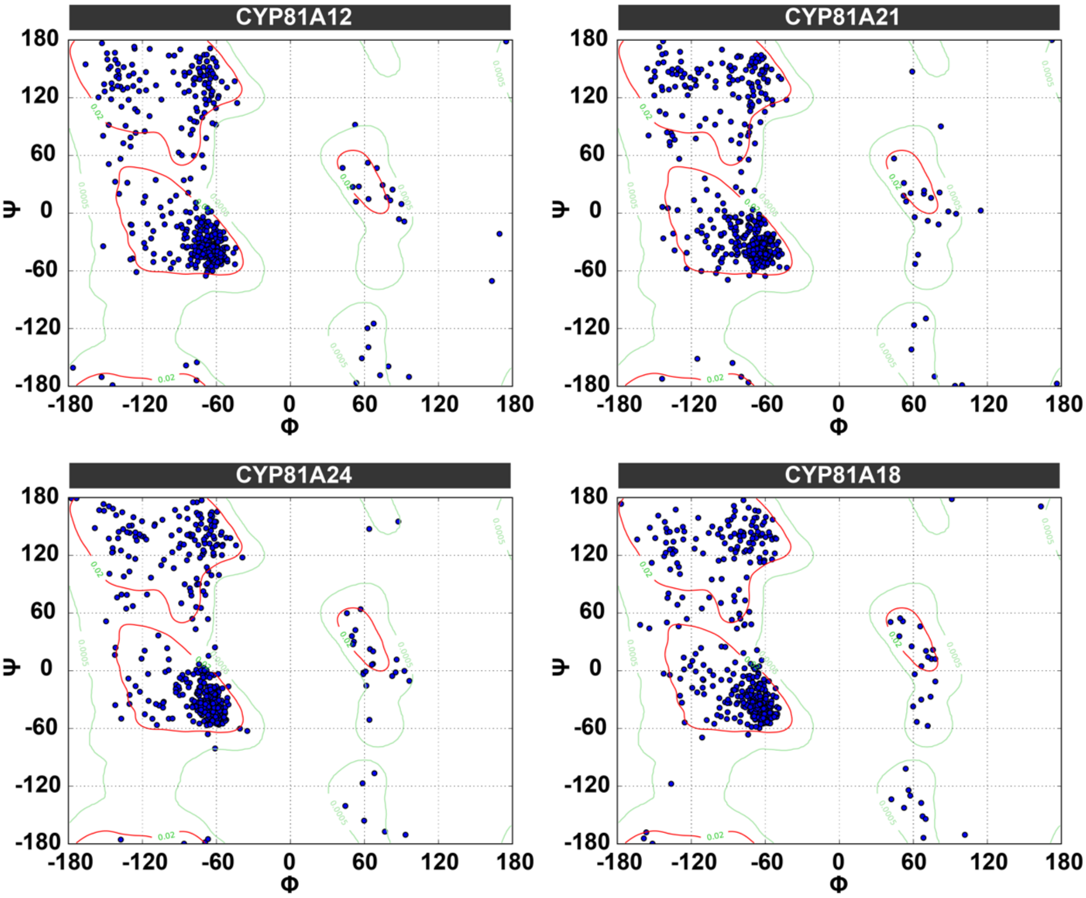


**Figure S2.** Ramachandran plots of ψ versus φ angles for all amino acids in CYP81A12, CYP81A21, CYP81A24, and CYP81A18, generated using RAMPAGE in UCSF Chimera. Blue dots represent individual residues, while red and green areas indicate favored and allowed regions, respectively, reflecting overall protein structure quality.
