## Supplementary material for "Promiscuous cytochrome P450s confer metabolic resistance to synthetic auxin herbicides in the weedy grass *Echinochloa phyllopogon*": Fig. S3

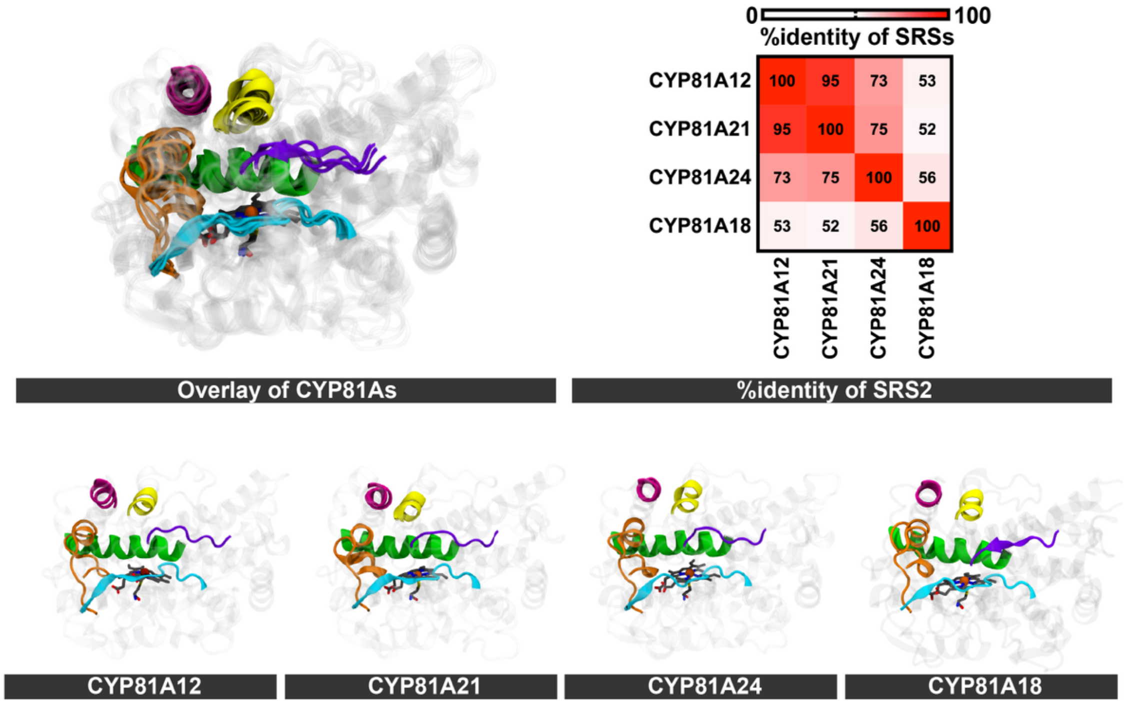


**Figure S3.** Structural models of CYP81A12, CYP81A21, CYP81A24, and CYP81A18 shown as both superimposed and individual structures. Pairwise structural alignment of substrate recognition sites (SRSs) highlights domain identity, visualized as a grid map with color intensity ranging from white (0%) to red (100%).
