## Supplementary material for "Promiscuous cytochrome P450s confer metabolic resistance to synthetic auxin herbicides in the weedy grass *Echinochloa phyllopogon*": Fig. S4

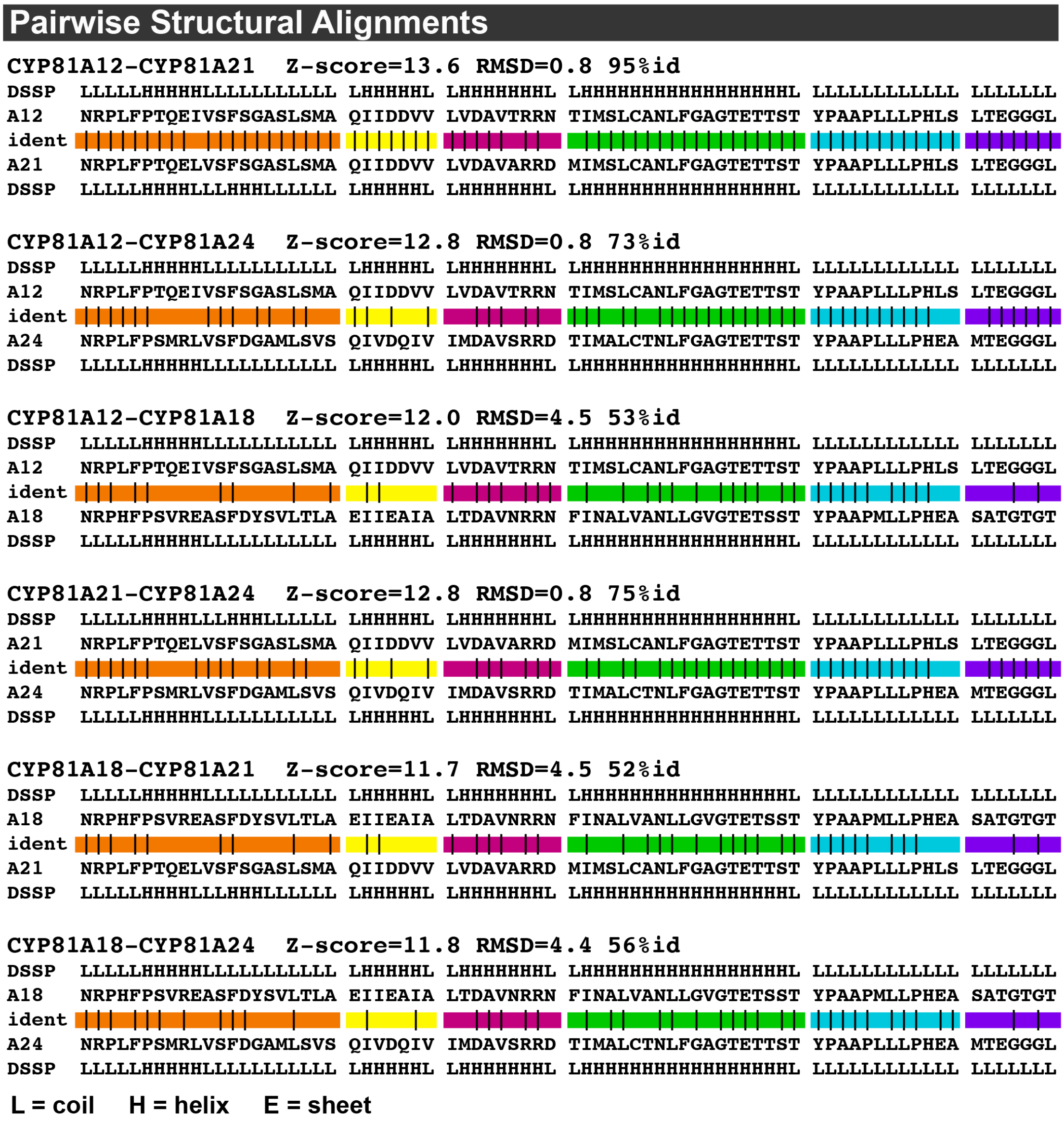


**Figure S4.** Pairwise structural alignment of SRS domains among CYP81A12, CYP81A21, CYP81A24, and CYP81A18. Structural identity between substrate recognition sites (SRSs) is shown based on pairwise comparisons of each CYP81A variant.
