## Supplementary material for "Promiscuous cytochrome P450s confer metabolic resistance to synthetic auxin herbicides in the weedy grass *Echinochloa phyllopogon*": Fig. S5

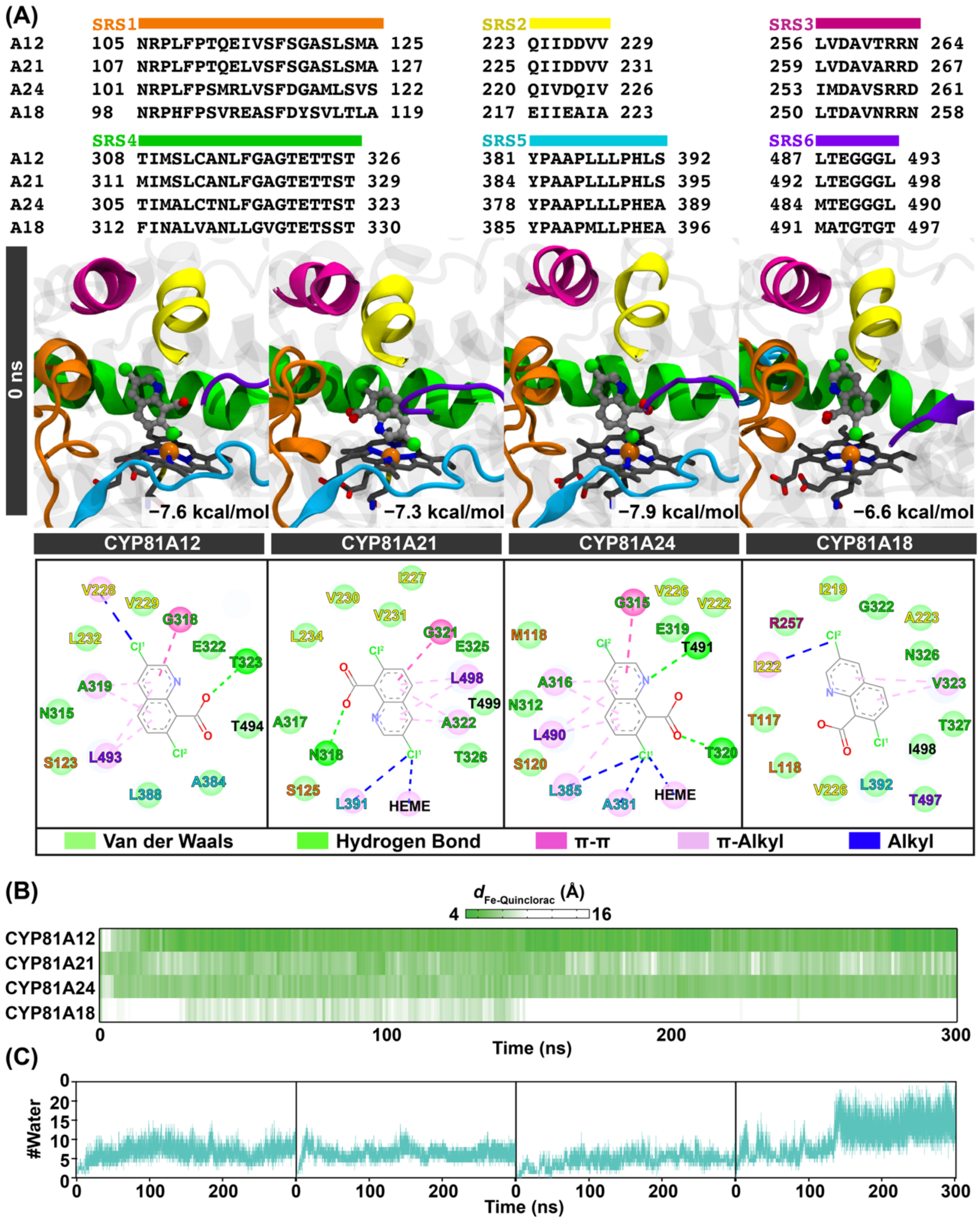


**Figure S5.** Structural and dynamic analysis of quinclorac binding in CYP81A variants. (A) SRS1–SRS6 sequence alignment, binding poses at 0 ns, and 2D interaction maps showing key residue contacts and $\Delta G_{bind}$ values. (B) Heatmap of quinclorac–heme distance ($d_{Fe\text{–}Quinclorac}$) over 300 ns MD trajectory. (C) Time evolution of water molecules in the binding pocket.
