## Supplementary material for "Promiscuous cytochrome P450s confer metabolic resistance to synthetic auxin herbicides in the weedy grass *Echinochloa phyllopogon*": Fig. S6

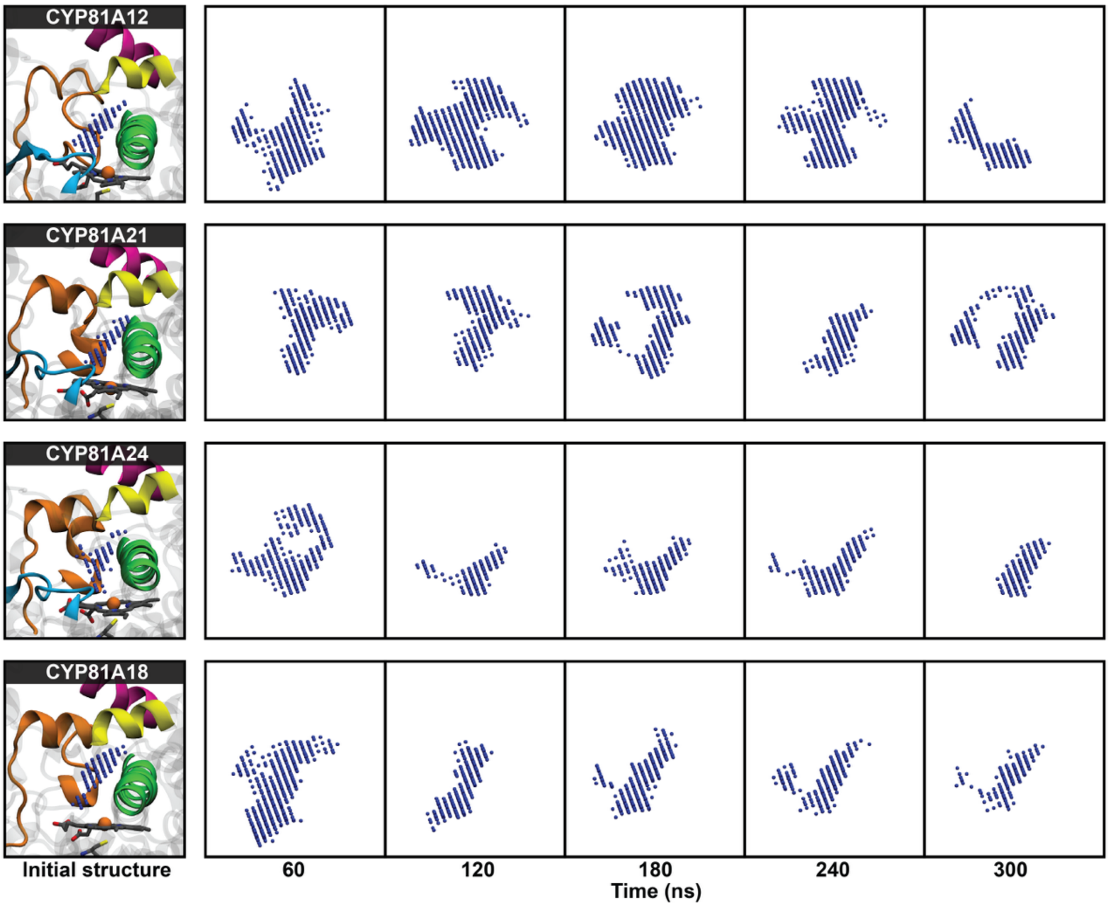


**Figure S6.** Volume analysis of CYP81A variant binding pockets over 300 ns MD simulations. The blue dots represent the volume of the CYP81A variant at each specific time point.
